# Multi-Tag: A modular platform of bioorthogonal probes for multi-modal (glyco)protein analysis

**DOI:** 10.1101/2022.10.24.513273

**Authors:** Marc D. Driessen, Hauke L. Junghans, Laura Hartmann, Ulla I. M. Gerling-Driessen

**Author notes:** These authors contributed equally (Marc D. Driessen, Hauke L. Junghans).

## Abstract

Bioorthogonal chemistry is a well-established concept for tagging and analyzing targets of interest even in living cells, tissue or organisms. In particular glycans, which are, as a posttranslational modification, not amenable to genetic engineering, became analytically accessible through the establishment of metabolic oligosaccharide engineering and subsequent bioorthogonal tagging of chemical probes. Since many essential cellular processes involve glycoproteins, it is not surprising that especially aberrant glycosylation has been associated with the pathology of many diseases. Investigation of aberrant glycosylation in a disease background is complicated by the heterogeneity of glycans and dynamic changes in their composition. In order to create a meaningful information depth, it can be beneficial to analyze the same sample with different analytical methods. This becomes even more relevant for samples with limited access. Most of the currently existing bioorthogonal probes are designed for use in only one type of experiment. These design restrictions are mainly based on the limited synthetic accessibility of more complex bioorthogonal probes. Multi-step syntheses are often time consuming and cost-inefficient. Here, we introduce a fast and easily manageable strategy for the synthesis of complex bioorthogonal probes that allow an application in multiple coordinated experiments. Using established principles and conditions of solid-phase peptide synthesis, we combine different functional building blocks to generate multi-functional bioorthogonal probes (named **Multi-Tag**s). We show the easy synthesis of several multi-modal probes and demonstrate their applicability and versatility in exemplary assays.

## INTRODUCTION

Protein glycosylation is one of the most prevalent posttranslational modifications (PTM) that substantially affects the structure and function of proteins in a wide variety of biological recognition events (intercellular interactions, pathogen recognition, immune modulation).^1-5^ Hence, aberrant glycosylation has been recognized to play a major role in various diseases.^6^ These range from genetic disorders with specific mutations of enzymes in the glycan biosynthesis pathways (summarized under the umbrella term **C**ongenital **D**isorders of **G**lycosylation; CDG)^7^ to dynamic environmental-induced changes that can be found in certain cancers^8-11^ or neurodegenerative diseases.^12-14^ One way to acquire a better understanding of cause and underlying pathology of these diseases, is the detailed mapping of the glycoproteome with combined information on proteins and the attached glycans.^15^ However, the investigation of intact glycoproteins remains challenging, as glycans are a non-templated PTM and as such not amenable by traditional genetic techniques. The heterogenic character of glycoproteins further complicates their analysis.^16^ Even available tools for untargeted analysis, such as MS-based proteomics, are hampered by significant signal dilution that result from the diversity of different glycoproteins isoforms existing in parallel.^17^ A given glycoprotein usually has more than one glycosite, that might or might not be occupied by a glycan (occupancy / macroheterogeneity) at a given time.^18^ Furthermore, the glycan structure occupying these sites can vary (microheterogeneity) leading to several isoforms.

One way to counteract some of these difficulties is the targeted chemical modification of proteins of interest. To this effect, bioorthogonal chemistry has long been utilized to covalently attach probes in a way that does not exhibit cross reactivity with the natural system.^19-22^ Metabolic incorporation of functionalized sugar analogs, so called metabolic oligosaccharide engineering (MOE),^23,24^ followed by tagging them with analytical tools has previously been established to analyze glycoproteins.^25,26^ To this end, bioorthogonal probes equipped with fluorophores or affinity tags have been developed to analyze glycans and glycoproteins in applications, such as live cell imaging^27^ or glycoproteomics.^28,29^ The latter chemoproteomics approach, a concept called Isotope Targeted Glycoproteomics (IsoTaG)^28,30^ is based on linking metabolically incorporated glycans derivatives to probes that contain heavy isotopes and biotin as an enrichment handle. The probe is designed to enrich tagged glycoproteins on a solid support (Streptavidin-coated agarose). These glycoproteins can be digested on-bead, keeping only the glycan modified peptides on the resin. The heavy isotopes recode the specific isotope pattern of the tagged glycopeptides, which can be used to specifically identify glycopeptide signals and allows for targeted tandem MS analysis of intact glycopeptides, even for low abundant species.^30^

Bioorthogonal probes have been established for a number of analytical methods over the years.^31^ However, most approaches are designed for one particular experiment or can be applied for one analytical read out (i.e., fluorophore labelling for imaging or attachment of an affinity tag for enrichment). Available probes are also often limited with regard to their bioorthogonal attachment chemistry (i.e., Cu-catalyzed azide-alkyne cycloaddition (CuAAC)^32^ vs. strain-promoted azide-alkyne cycloaddition (SPAAC)^33^. For example, in some cases, probes might be available with an azide or alkyne but not with a DBCO or TCO group. This can make a multiplex or multi-use application very difficult and expensive. Another limitation of more complex probes that find, for instance, application in chemoproteomics, is their synthesis, which often involve complex synthetic and purification procedures, such as inert conditions and column chromatography. Such multi-step synthesis protocols are usually time-consuming and may give rise to low overall yields.

To overcome this limitation, we developed a strategy for an easy and fast generation of bioorthogonal probes that allows the combination of multiple experiments by using only one tag that is attached to the sample of interest. These probes can carry any bioorthogonal handle on one end and can be equipped with multiple functional units in a building block fashion (e.g., fluorophores, cross linkers, cleavable linkers, affinity tags etc.) on the other end, to allow a combination of different analytical read out options from one sample. The synthesis of these probes is based on combining the building blocks via the well-established solid-phase peptide synthesis (SPPS) protocols.^34^ The modular strategy offers free choice of combining the individual functional units which allows the combination of multiple experimental readouts with one probe in smart workflows. Our strategy further allows the introduction of any bioorthogonal group to every probe thereby offering a huge flexibility with regard to the bioorthogonal attachment strategy that can be applied with these probes.^35^ Here, we present our SPPS-based synthesis strategy and the first set of such multi-functional bioorthogonal probes. The presented strategy allows a great flexibility for customizing the probes according to the individual demands of the experiments. We aim to apply our **multi-**functional probes for the **tag**ging and analysis of metabolically engineered (glyco)proteins (**Multi-Tag**).

## RESULTS AND DISCUSSION

### Probe design and synthesis

The Multi-Tag concept is based on the idea to gain easy access to complex biorthogonal probes that can be used in multiple experiments. This requires the combination of multiple functionalities (here referred to as functional units, *Figure 1*), such as fluorophores, cleavable linkers, affinity tags etc. in one probe. Aiming for a system that can provide high flexibility with regard to the combination of these individual functional units, we identified solid-phase peptide synthesis as an ideal strategy for this purpose (*Figure 1A*). We selected building blocks that, similar to amino acids, contained one carboxy group and one primary amine (*Figure 1B*) and combined them using previously described standard 9-Fluorenylmethoxycarbonyl (Fmoc)-strategy with Benzotriazole-1-yl-oxy-tris-pyrrolidino-phosphonium hexafluorophosphate (PyBOP) and N, N-Diisopropylethylamin (DIPEA) as coupling reagents. Given the modularity of the SPPS approach, we realized the synthesis of a small library of multi-functional bioorthogonal probes that vary in complexity due to the number of functional units that were combined per probe (*Figure 1*).

**Figure 1:**
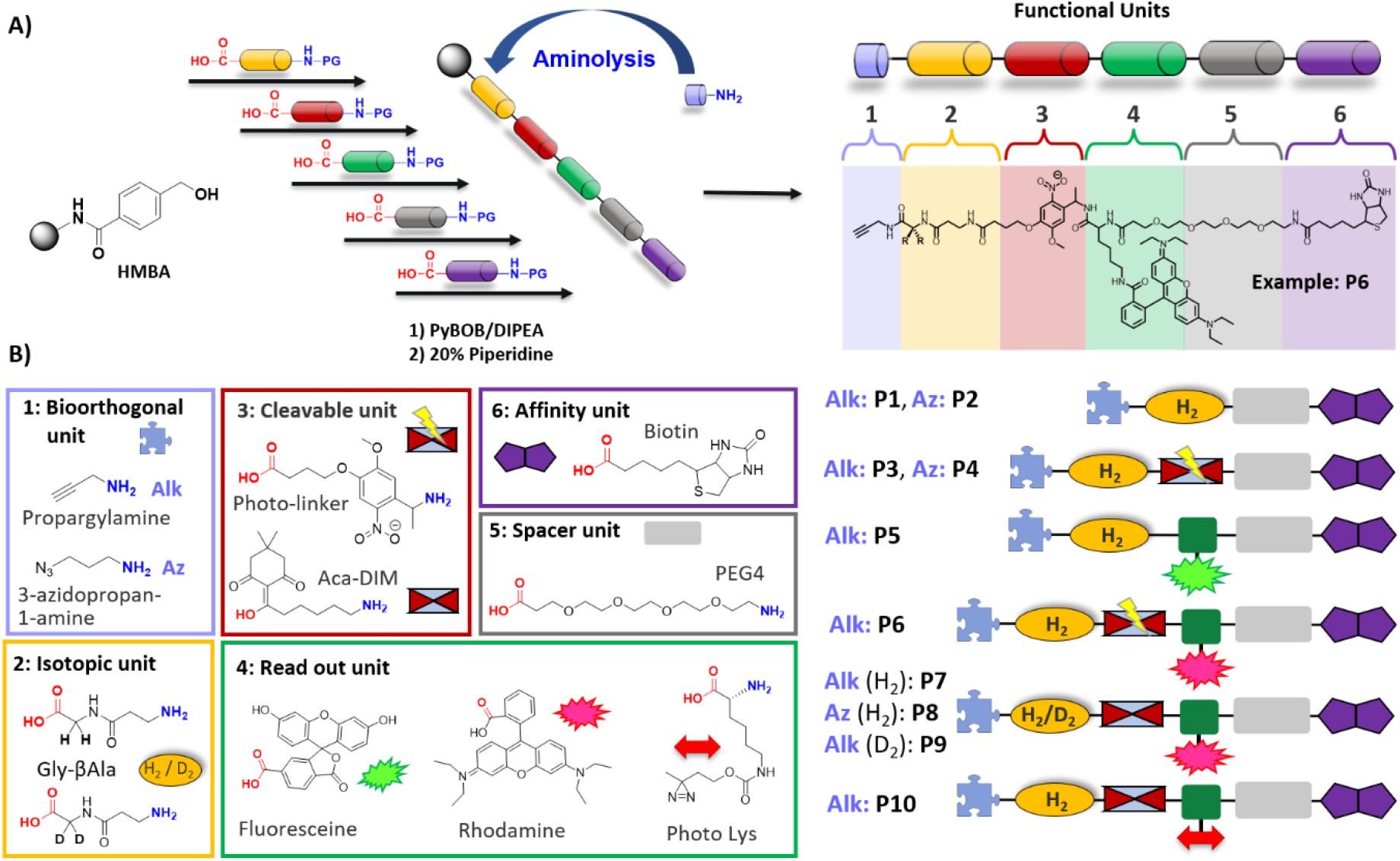
**A)** Schematic representation of SPPS-based synthesis of multi-functional bioorthogonal probes. **B)** Functional building blocks selected to generate the first generation of Multi-Tags (P1-P10).

The design of this first generation of probes was chosen to allow a combination of either imaging or proximity labelling experiment with subsequent enrichment of the tagged sample and further analysis via Mass spectrometry. Thus, all probes contain an N-terminal PEG_4_-Biotin as the affinity unit, which allows enrichment and isolation of attached samples based on binding to streptavidin-coated beads. Inspired by the Iso-TaG approach,^14^ which uses heavy isotopes to recode the isotopic pattern of tagged proteins, we wanted to include this feature in our probe design. Hence, all probes presented here, contain a dipeptide core structure consisting of glycine and β-alanine. Glycine is available in different isotopic versions (e.g., D_2_, ^13^C, ^15^N), which provides the opportunity to equip every probe with an inherent label to be utilized either in tandem MS-experiments similar to the IsoTaG^14^ concept or for MS-based quantification of tagged proteins.^36^ The other functional units were varied in the individual probes presented here. The most complex designs of this first generation of multi-functional probes combine 6 functional units. The probes (P6 – P10) contain an azide or alkyne as **bioorthogonal** handle, the **isotopic core** unit (with either Gly or deuterated Gly), a **cleavable linker** next to a branched building block (lysine) carrying a **fluorophore** (or a **photo crosslinker**) and PEG_4_-**spacer**-bound biotin as **affinity tag** (*Figure 1B*).

### Introducing the bioorthogonal handle of choice

Our second goal was to achieve flexibility with regard to the bioorthogonal handle that the probes can be equipped with. Therefore, we used a 4-(Hydroxmethyl)benzoic acid (HMBA)-linker on the resin, which can be cleaved under basic or nucleophilic conditions using O-, N- or S-nucleophiles.^37^ Using amines that carry a functional group of choice, probe release from the solid support and C-terminal introduction of the functional group can be performed in one step. We chose amines that carry a bioorthogonal groups (e.g., propargylamine or 3-azidopropan-1-amine) to introduce them during the final step of the synthesis. In agreement with previous findings,^37^ we see high yields of C-terminal functionalization for amines with low steric demands, such as propargylamine or 3-azidopropan-1-amine (*Figure 2A*). While cleavage and C-terminal functionalization can be detected one hour after addition of the respective amines, full conversion requires longer reaction times of up to 20 hours, as has been previously reported. ^37^ For amines carrying functional groups with higher steric demand, such as DBCO or tetrazine, we observed only low conversion if they were used as single reagents to directly cleave the probe off the resin. In order to also introduce these sterically demanding bioorthogonal handles, we adapted a protocol from Hickey et al.,^38^ where the authors showed the synthesis of cyclic peptides by cleaving the HMBA-linker with 1,5,7-triazabicyclo[4.4.0]dec-5-en (TBD), which forms a C-terminal active ester upon linker hydrolysis that subsequently reacts with the free amine on the N-terminus of the peptide strand. Based on above-described design principle, our probes contain an N-terminal biotin. Lacking any free amines, these probes do not undergo intramolecular cyclization. Thus, we use TBD to first cleave the HMBA-linker and form a C-terminal TBD-active ester, which then reacts with the functionalized amine that is added to the reaction mixture to give the bioorthogonal handle on the C-terminus (*Figure 2B*). Conversions can be further increased by adding equimolar amounts of PyBOP/ DIPEA to the reaction mixture but require a purification step afterwards. Deviating from the reported protocol,^38^ we observed less side products formation using DCM instead of DMF as solvent for this step.

**Figure 2:**
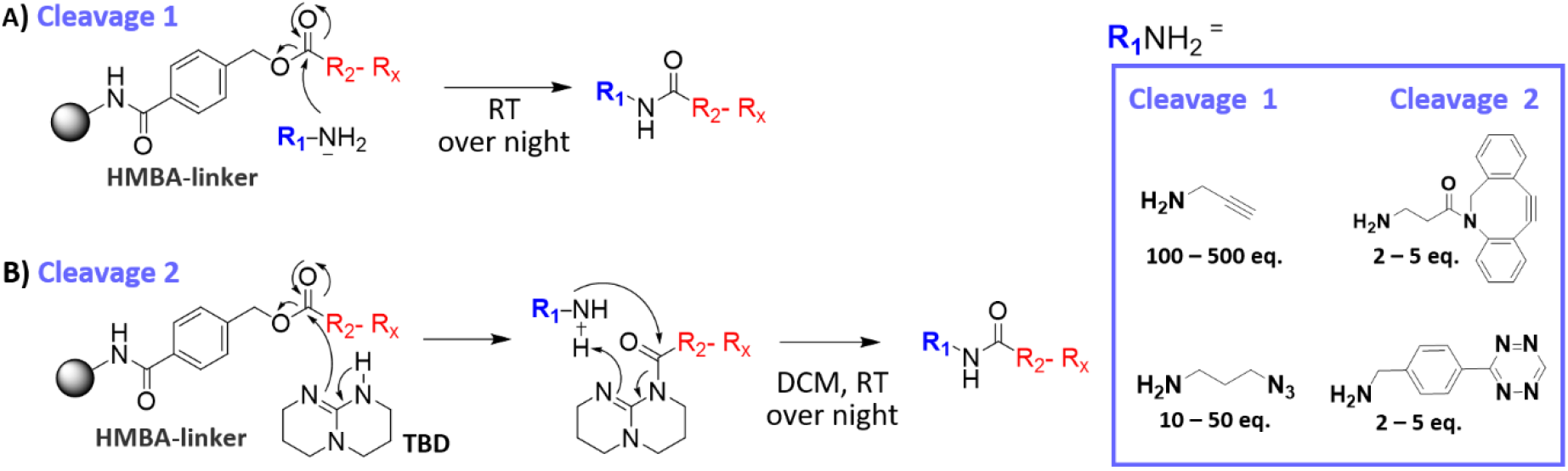
**A)** Nucleophilic cleavage of the HMBA-linker with functionalized amines to generate C-terminal functionalized probes. **B)** Cleavage of the HMBA-linker using TBD and subsequent C-terminal probe functionalization.

The SPPS concept allows batch synthesis of the multi-functional probes and storage on the solid support. Each probe can be equipped with a variety of bioorthogonal handles by simply dividing the resin into multiple portions prior to cleavage and C-terminal functionalization. This provides high flexibility of the presented probes with regard to the labeling strategy of the targets (e.g., Cu-catalyzed 1,3-dipolar cycloaddition (CuAAC), Strain-promoted azide alkyne cycloaddition (SPAAC) or inverse electron demanding Diels–Alder reaction (IEDDA)^39,40^ and allows a combined use of multiple probes in one system applying orthogonal bioorthogonal reactions.

### Sample tagging and analysis

Using a representative set of probes with affinity labels and various fluorophores, we performed experiments to demonstrate their usability and versatility.

In a first experiment, we used a serial dilution set up to estimate detection limits. We introduced Azides into BSA via non-specific HNS-based coupling to lysine side chains. We used the Azide-modified BSA and tagged it with **P7** using CuAAC and then diluted it several times by a factor of 5 in a full cell lysate (LN18, a glioblastoma cell line). Starting with 20μg of P7@**BSA-N**_3_ in 20 μg of lysate, we made 7 dilutions of 4μg/0.8μg/160ng/32ng/6.4ng/1.3ng/0.26ng respectively. **P7** contains Rhodamine as fluorophore, DDE as a hydrazin-cleavable linker and biotin as an affinity label. To analyze (**P7@BSA-N**_**3**_), we detected in-gel fluorescence of the rhodamine dye and streptavidin-HRP bound to the biotin moiety of P7 after transferring the same gel to a nitrocellulose membrane (western blot). Rhodamine signals in the gel are visible, albeit very faint down to between 800ng-160ng, while biotin/streptavidin-HRP chemiluminescence signals in the western blots are visible down to the lower ng dilution upon long detection times (Figure 3). Similar results were achieved when first diluting BSA-N_3_ and then performing the click reaction (data not shown). The lysate contained several strong biotinylated background bands directly above the **P7@BSA-N**_**3**_ band, yet the bands could be clearly distinguished. Total protein stains of the same blot showed no discernable differences between the lanes. These results were satisfying in itself, as a reasonably low detection limit, both in-gel and by WB-detection could be established. During sample preparation, we noticed that fluorescence could already be detected in the vessels used for dilution and thought that this could be used as a useful indicator for successful tagging of Multi-Tags that contain fluorophores.

**Figure 3:**
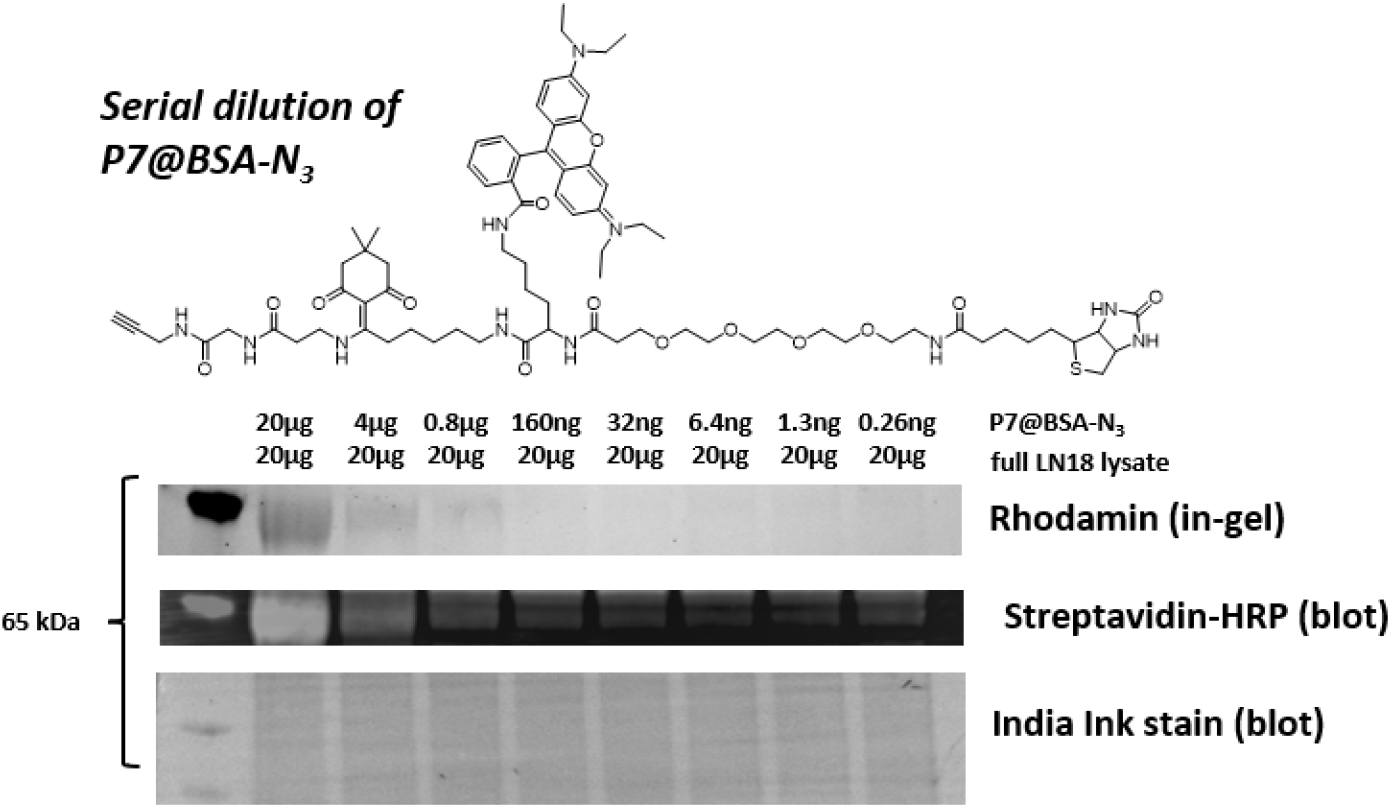
Detection of **P7** tagged to azide-modified BSA in different dilutions in complex cell lysate. The rhodamine label was detected by in-gel fluorescence and the biotin label by chemiluminescence after incubating the western blot with HRP-labeled streptavidin. Indira Ink stain was used to visualize the total protein content per lane.

### Multi-tag allows optical workflow control at every step

In the next step, we used a stronger fluorescing probe **P5** carrying a fluorescein-based dye as well as biotin, again conjugated to the same model protein to give **P5@BSA-N**_**3**_. Here, we performed a 10x serial dilution of **P5**@**BSA-N**_**3**_ in the same lysate (LN18). Starting with 4μg of **BSA-N**_**3**_ in 20μg LN18 lysate gave dilutions of 0.4μg/40ng/4ng/0.4ng/40pg/4pg/0.4pg of **P5**@**BSA-N**_**3**_ (*Figure 4*). We first determined fluorescence directly in the PCR-tubes that contained the dilution series and were used to load on the gel (clear signal down to 400ng). We then detected fluorescence in the SDS-PAGE gel directly after electrophoresis (signal visible down to 40 ng) and this time also on the corresponding western blot, yielding (faint) signals down to 0.4ng-4 ng). The subsequent incubation with streptavidin-HRP allowed a detection of **P5@BSA-N**_**3**_ down to approximately 0.4 ng and below with long exposure times. Total protein stain again showed no visible loading differences of the samples in the individual lanes, showing that the lysate is the main determinant of sample composition. This workflow shows one of the big advantages garnered through the use of multi-functional probes: At every experimental step an immediate, quick and loss-free quality control of the workflow can be recorded. The workflow presented here lends itself to every step and type of experiment:

**Figure 4:**
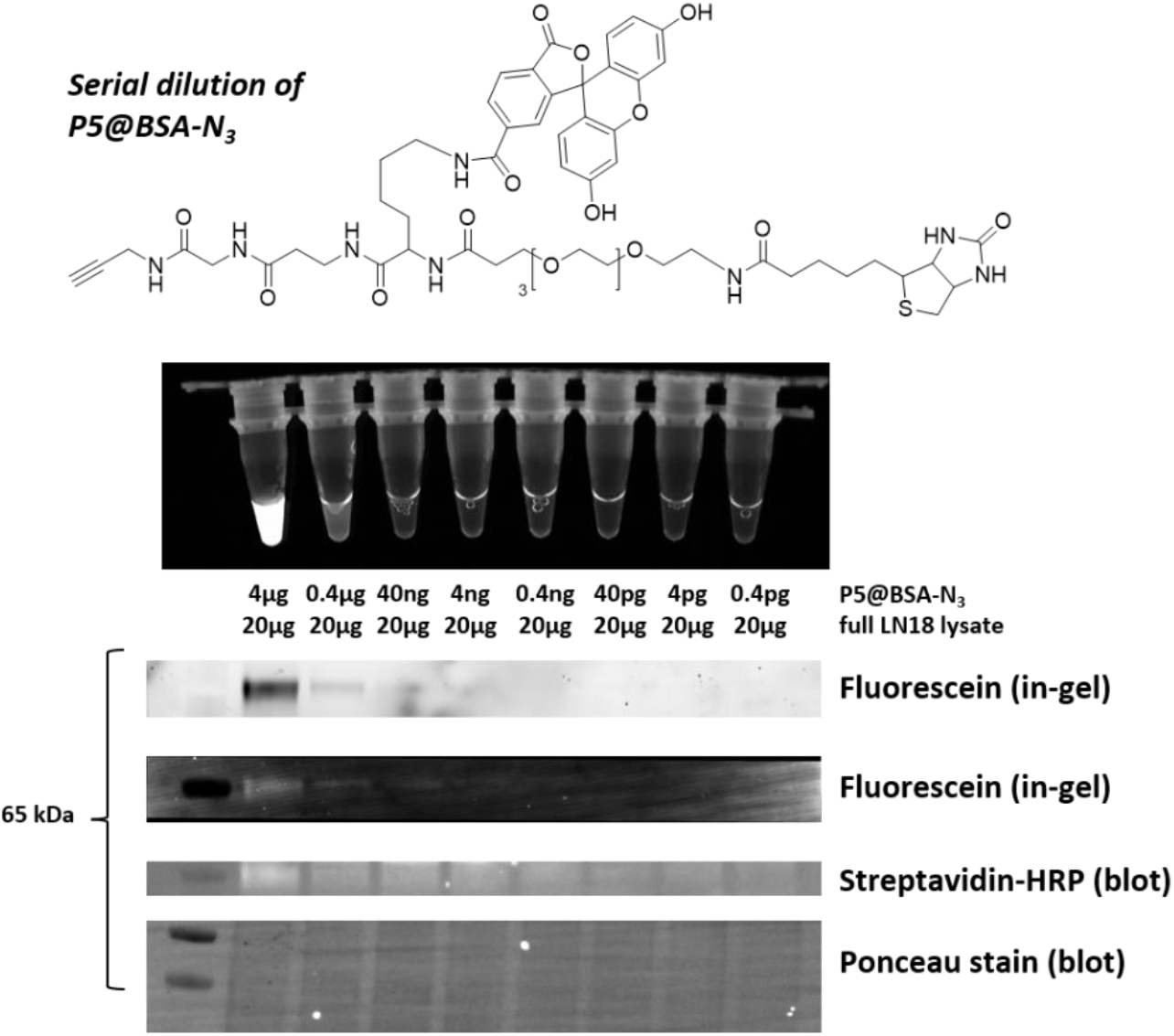
Detection of **P5** tagged to azide-modified BSA in different dilutions in complex cell lysate. The fluorescence of Fluoresceine was detected directly in the sample tubes, in the SDS-PAGE gel and on the western blot. The biotin label was visualized chemiluminescence after incubation of the western blot with HRP-labeled streptavidin. Indira Ink stain was used to visualize the total protein content per lane.

1. Click-modification of targets followed by precipitation – optical control “in-tube”.
2. Enrichment – optical control of beads/supernatant “in-tube”
3. SDS-PAGE separation – in-gel control
4. Wester blotting – on-blot control

However, it has to be noted that using the SDS-PAGE for detection is less sensitive, if the gel is to be used for blotting – as fixing and thorough washing of the gel is not possible, interference and dispersion of sample make it less sensitive than a thoroughly fixed and washed gel. In addition, if the detection steps take too long, signal loss due to photo bleaching and washing out of the sample can be observed in western blotting.

### Multi-Tag labeling of enzymatically labelled HEK cells

To test Multi-Tag in complex samples, we used the commercial click-it O-GlcNAc kit by Thermo Fischer to label a full lysate of HEK cells. Identical “click” reaction of **P6** with labelled and unlabeled lysates (*Figure 5*) were performed in parallel. We detected Rhodamine fluorescence in gel and after blotting and then used AF488 labeled streptavidin to co-detect fluorescein and biotin signals. Even though the resolution of the labelled lysates was not ideal, it clearly showed difference in signal intensity of the Rhodamine signal as well as a colocalization with the biotin signals, which indicates that the Multi-Tag only attached to cell lysate containing the enzymatically labelled GalNAz.

**Figure 5:**
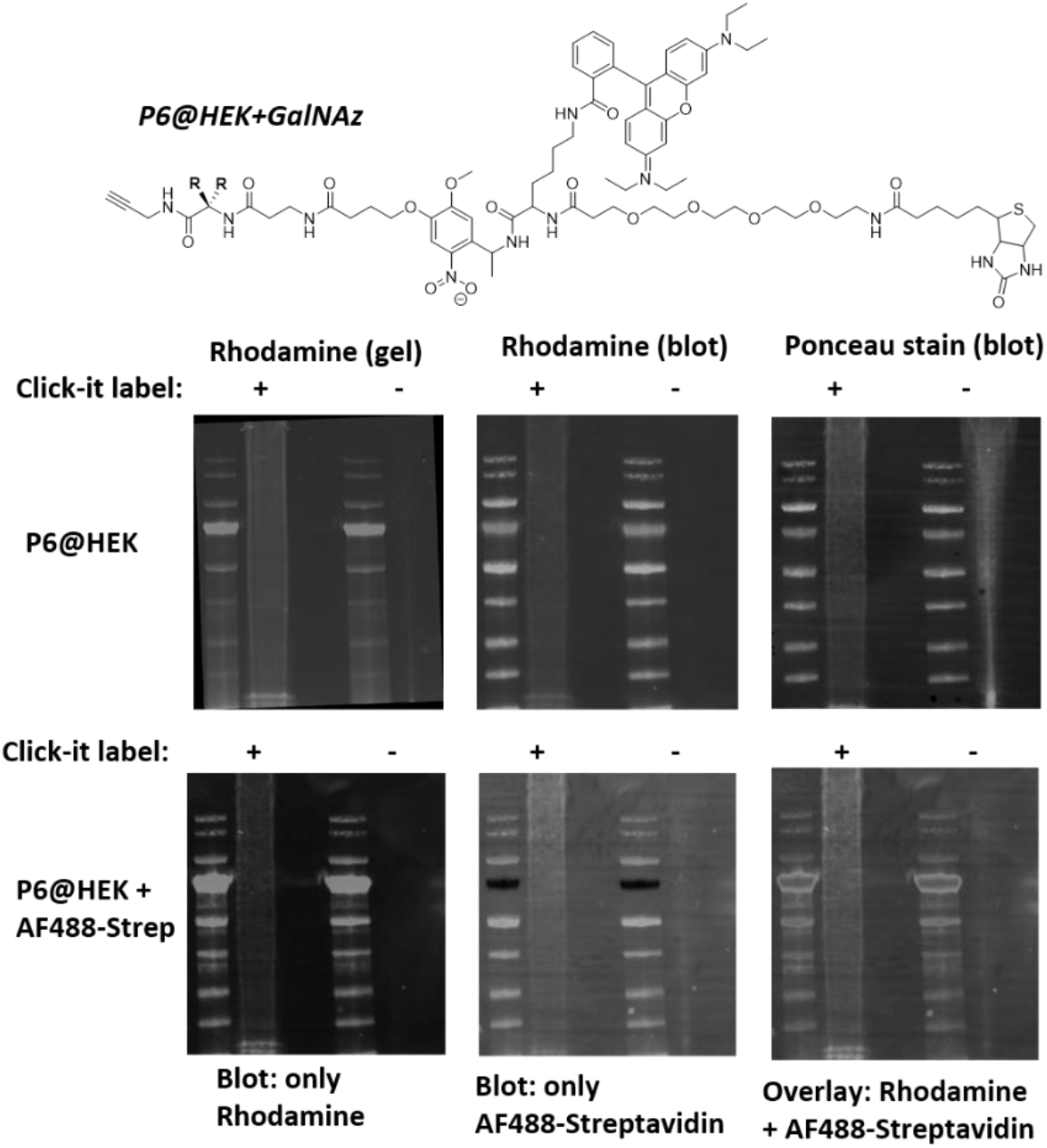
Detection of **P6** tagged to enzymatically GalNAz labelled cell lysate by visualization of the Rhodamine fluorescence in gel and on the western blot. The biotin label was visualized chemiluminescence after incubation of the western blot with HRP-labeled streptavidin. Ponceau stain was used to visualize the total protein content per lane.

## CONCLUSION

In summary, we present a fast and easy strategy based on standard solid-phase synthesis procedures to give access to complex bioorthogonal probes (here called Multi-Tags) that allow the analysis of tagged samples with multiple analytical methods in coordinated workflows. The probes consist of various functional units that are combined in a highly-flexible modular fashion using established coupling and deprotection protocols. Using an HMBA-linker, which can be cleaved with functional nucleophiles (here primary amines) allows the C-terminal functionalization during the final step of the synthesis. This provides further flexibility with regard to the bioorthogonal groups that the probes can be equipped with. The highly efficient SPPS approach does not require time-consuming purification steps. Final products can be purified on a single solid-phase extraction column using C18-functionalized silica as the stationary phase material. Multi-Tags of the first generation presented here, allow, as an example of coordinated experiments, a combination of imaging or proximity labelling with sample enrichment and further MS-analysis. However, the modular design of this platform combined with the easy synthetic access allows the combination of many other potential functional units (*Figure 6*) to create probes that can, based on used building blocks,^41-43^ be modulated with regard to their physicochemical properties and customized to the desired experimental workflow. We initially established Multi-tags for the analysis of bioorthogonal engineered glycoproteins but we believe that the modular platform holds great potential for the use in many other applications.

**Figure 6:**
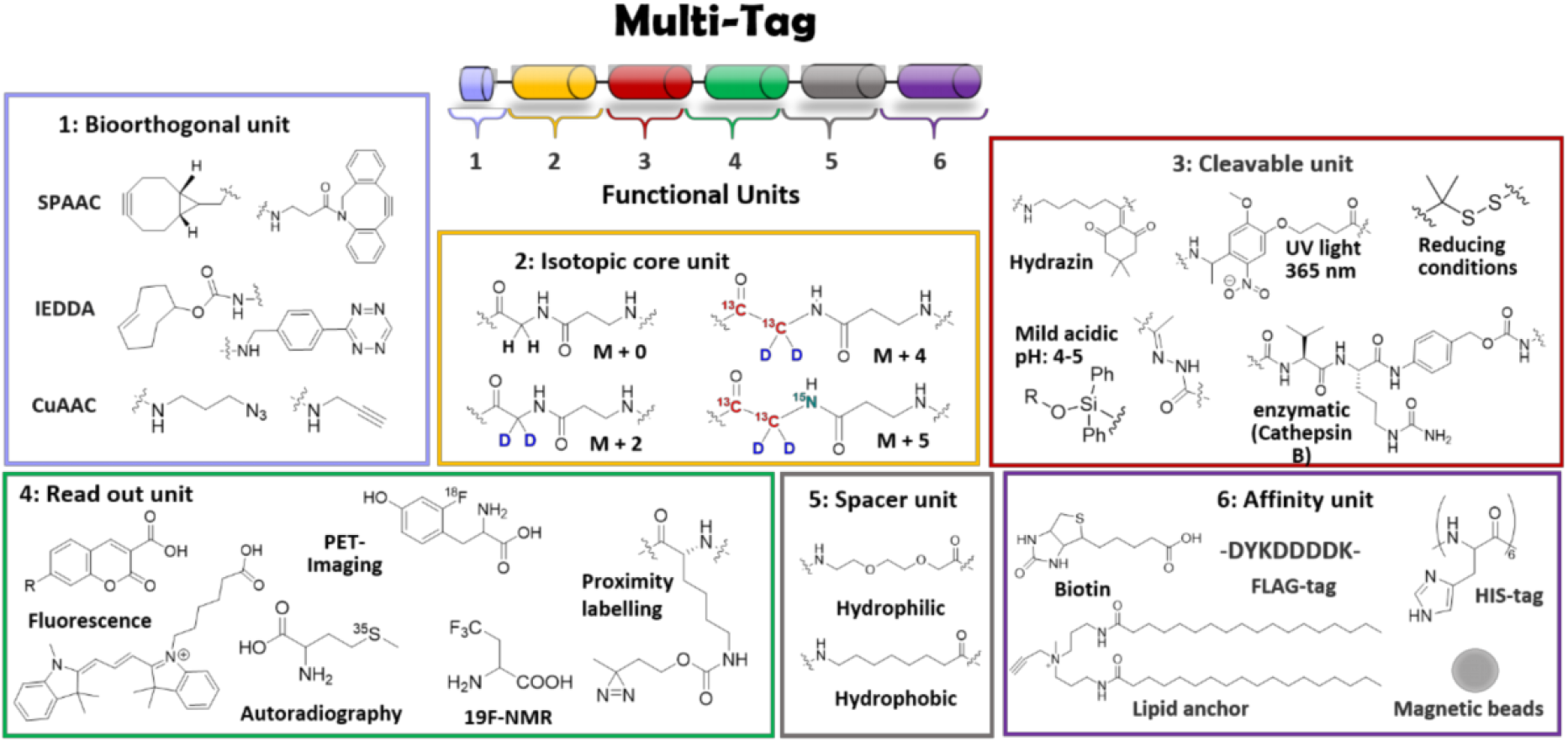
Selection of additional potential functional units that are compatible with the SPPS-based synthesis strategy of the Multi-Tag platform.

## Acknowledgements

We thank Professor Dr. Carolyn Bertozzi for foundational discussions. U. G.-D. is supported through funding from the SFF - Strategic Research Fund of the Heinrich Heine University Düsseldorf (Project GlycoPathogens). M.D.D. is funded by the Deutsche Krebshilfe, through a Mildred Scheel Nachwuchszentrum Grant (Grant number 70113307).

## References

(1) Varki, A. Biological roles of glycans. Glycobiology 2016, 27 (1), 3.

(2) Lin, B.; Qing, X.; Liao, J.; Zhuo, K. Role of Protein Glycosylation in Host-Pathogen Interaction. Cells 2020, 9, 1022.

(3) Amon, R.; Reuven, E. M.; Leviatan Ben-Arye, S.; Padler-Karavani, V. Glycans in immune recognition and response. Carbohydr. Res. 2014, 389, 115.

(4) Grivennikov, S. I.; Greten, F. R.; Karin, M. Immunity, Inflammation, and Cancer. Cell 2010, 140, 883.

(5) Heine, V.; Hovorková, M.; Vlachová, M.; Filipová, M.; Bumba, L.; Janoušková, O.; Hubálek, M.; Cvačka, J.; Petrásková, L.; Pelantová, H. et al. Immunoprotective neo-glycoproteins: Chemoenzymatic synthesis of multivalent glycomimetics for inhibition of cancer-related galectin-3. Eur. J. Med. Chem. 2021, 220, 113500.

(6) Reily, C.; Stewart, T. J.; Renfrow, M. B.; Novak, J. Glycosylation in health and disease. Nat. Rev. Nephrology 2019, 15 (1), 346.

(7) Péanne, R.; de Lonlay, P.; Foulquier, F.; Kornak, U.; Lefeber, D. J.; Morava, E.; Pérez, B.; Seta, N.; Thiel, C.; Van Schaftingen, E. et al. Congenital disorders of glycosylation (CDG): Quo vadis? Eur. J. Med. Genet. 2018, 61 (1), 643.

(8) Pinho, S. S.; Reis, C. A. Glycosylation in cancer: mechanisms and clinical implications : Nature Reviews Cancer : Nature Publishing Group. Nat. Rev. Cancer 2015, 15, 540.

(9) Stowell, S. R.; Ju, T.; Cummings, R. D. Protein Glycosylation in Cancer. Annu. Rev. Pathol. 2015, 10, 473.

(10) Oliveira-Ferrer, L.; Legler, K.; Milde-Langosch, K. Role of protein glycosylation in cancer metastasis. Semin. Cancer Biol. 2017, 44, 141.

(11) Dube, D. H.; Bertozzi, C. R. Glycans in cancer and inflammation — potential for therapeutics and diagnostics. Nat. Rev. Drug Discov. 2005, 4, 477.

(12) Botella-López, A.; Burgaya, F.; Gavín, R.; García-Ayllón, M. S.; Gómez-Tortosa, E.; Peña-Casanova, J.; Ureña, J. M.; Del Río, J. A.; Blesa, R.; Soriano, E. Reelin expression and glycosylation patterns are altered in Alzheimer’s disease. Proc. Natl. Acad. Sci. U.S.A. 2006, 103 (14), 5573.

(13) Wang, A. C.; Jensen, E. H.; Rexach, J. E.; Vinters, H. V.; Hsieh-Wilson, L. C. Loss of O-GlcNAc glycosylation in forebrain excitatory neurons induces neurodegeneration. Proc. Natl. Acad. Sci. U.S.A. 2016, 113 (1), 15120.

(14) Moll, T.; Shaw, P. J.; Cooper-Knock, J. Disrupted glycosylation of lipids and proteins is a cause of neurodegeneration. Brain 2020, 143 (1), 1332.

(15) Chang, I. J.; He, M.; Lam, C. T. Congenital disorders of glycosylation. Annals of translational medicine 2018, 6 (1), 477.

(16) Rudd, P. M.; Dwek, R. A. Glycosylation: Heterogeneity and the 3D Structure of Proteins. Crit. Rev. Biochem. Mol. Biol. 1997, 32, 1.

(17) Bagdonaite, I.; Malaker, S. A.; Polasky, D. A.; Riley, N. M.; Schjoldager, K.; Vakhrushev, S. Y.; Halim, A.; Aoki-Kinoshita, K. F.; Nesvizhskii, A. I.; Bertozzi, C. R. et al. Glycoproteomics. Nat. Rev. Methods Primers 2022, 2 (1), 48.

(18) čaval, T.; Heck, A. J. R.; Reiding, K. R. Meta-heterogeneity: Evaluating and Describing the Diversity in Glycosylation Between Sites on the Same Glycoprotein. Mol. Cell. Proteomics 2021, 20.

(19) Ramil, C. P.; Lin, Q. Bioorthogonal chemistry: strategies and recent development. Chem. Commun. (Camb.) 2013, 49, 11007.

(20) Sletten, E. M.; Bertozzi, C. R. Bioorthogonal Chemistry: Fishing for Selectivity in a Sea of Functionality. Angew. Chem. Int. Ed. 2009, 48, 6974.

(21) Sletten, E. M.; Bertozzi, C. R. From mechanism to mouse: a tale of two bioorthogonal reactions. Acc. Chem. Res. 2011, 44, 666.

(22) Bertozzi, C. R. A Decade of Bioorthogonal Chemistry. Acc. Chem. Res. 2011, 44, 651.

(23) Gilormini, P.-A.; Batt, A. R.; Pratt, M. R.; Biot, C. Asking more from metabolic oligosaccharide engineering. Chem. Sci. 2018, 9 (1), 7585.

(24) Cioce, A.; Bineva-Todd, G.; Agbay, A. J.; Choi, J.; Wood, T. M.; Debets, M. F.; Browne, W. M.; Douglas, H. L.; Roustan, C.; Tastan, O. Y. et al. Optimization of Metabolic Oligosaccharide Engineering with Ac4GalNAlk and Ac4GlcNAlk by an Engineered Pyrophosphorylase. ACS Chem. Biol. 2021, 16 (1), 1961.

(25) Robinson, P. V.; de Almeida-Escobedo, G.; de Groot, A. E.; McKechnie, J. L.; Bertozzi, C. R. Live-Cell Labeling of Specific Protein Glycoforms by Proximity-Enhanced Bioorthogonal Ligation. J. Am. Chem. Soc. 2015, 137, 10452.

(26) Saxon, E.; Bertozzi, C. R. Cell Surface Engineering by a Modified Staudinger Reaction. Science 2000, 287 (1), 2007.

(27) Agarwal, P.; Beahm, B. J.; Shieh, P.; Bertozzi, C. R. Systemic Fluorescence Imaging of Zebrafish Glycans with Bioorthogonal Chemistry. Angew. Chem. 2015, 54, 11504.

(28) Woo, C. M.; Iavarone, A. T.; Spiciarich, D. R.; Palaniappan, K. K.; Bertozzi, C. R. Isotope-targeted glycoproteomics (IsoTaG): A mass-independent platform for intact N- and O-glycopeptide discovery and analysis. Nat. Methods 2015, 12, 561.

(29) Calle, B.; Bineva-Todd, G.; Marchesi, A.; Flynn, H.; Ghirardello, M.; Tastan, O. Y.; Roustan, C.; Choi, J.; Galan, M. C.; Schumann, B. et al. Benefits of Chemical Sugar Modifications Introduced by Click Chemistry for Glycoproteomic Analyses. J. Am. Soc. Mass. Spectrom. 2021, 32 (1), 2366.

(30) Woo, C. M.; Felix, A.; Byrd, W. E.; Zuegel, D. K.; Ishihara, M.; Azadi, P.; Iavarone, A. T.; Pitteri, S. J.; Bertozzi, C. R. Development of IsoTaG, a Chemical Glycoproteomics Technique for Profiling Intact N- and O-Glycopeptides from Whole Cell Proteomes. J. Proteome Res. 2017, 16, 1706.

(31) Devaraj, N. K. The Future of Bioorthogonal Chemistry. ACS Cent. Sci. 2018, 4, 952.

(32) Meldal, M.; Diness, F. Recent Fascinating Aspects of the CuAAC Click Reaction. Trends in Chemistry 2020, 2 (1), 569.

(33) Agard, N. J.; Prescher, J. A.; Bertozzi, C. R. A Strain-Promoted [3 + 2] Azide–Alkyne Cycloaddition for Covalent Modification of Biomolecules in Living Systems. J. Am. Chem. Soc. 2004, 126 (1), 15046.

(34) Palomo, J. M. Solid-phase peptide synthesis: an overview focused on the preparation of biologically relevant peptides. RSC Advances 2014, 4 (1), 32658.

(35) Nguyen, S. S.; Prescher, J. A. Developing bioorthogonal probes to span a spectrum of reactivities. Nat. Rev. Chem. 2020, 4, 476.

(36) Chahrour, O.; Cobice, D.; Malone, J. Stable isotope labelling methods in mass spectrometry-based quantitative proteomics. J. Pharm. Biomed. Anal. 2015, 113, 2.

(37) Hansen, J.; Diness, F.; Meldal, M. C-Terminally modified peptides via cleavage of the HMBA linker by O-, N-or S-nucleophiles. Org. Biomol. Chem. 2016, 14 (1), 3238.

(38) Hickey, J. L.; Lin, S. One-pot peptide cleavage and macrocyclization through direct amidation using triazabicyclodecene. Pept. Sci. 2020, 112 (1), e24161.

(39) Blackman, M. L.; Royzen, M.; Fox, J. M. Tetrazine ligation: fast bioconjugation based on inverse-electron-demand Diels-Alder reactivity. J. Am. Chem. Soc. 2008, 130, 13518.

(40) Dommerholt, J.; van Rooijen, O.; Borrmann, A.; Guerra, C. F.; Bickelhaupt, F. M.; van Delft, F. L. Highly accelerated inverse electron-demand cycloaddition of electron-deficient azides with aliphatic cyclooctynes. Nat. Commun. 2014, 5, 5378.

(41) Ponader, D.; Wojcik, F.; Beceren-Braun, F.; Dernedde, J.; Hartmann, L. Sequence-Defined Glycopolymer Segments Presenting Mannose: Synthesis and Lectin Binding Affinity. Biomacromolecules 2012, 13 (1), 1845.

(42) Boden, S.; Reise, F.; Kania, J.; Lindhorst, T. K.; Hartmann, L. Sequence-Defined Introduction of Hydrophobic Motifs and Effects in Lectin Binding of Precision Glycomacromolecules. Macromol. Biosci. 2019, 19 (1), 1800425.

(43) Morstein, J.; Capecchi, A.; Hinnah, K.; Park, B.; Petit-Jacques, J.; Van Lehn, R. C.; Reymond, J.-L.; Trauner, D. Medium-Chain Lipid Conjugation Facilitates Cell-Permeability and Bioactivity. J. Am. Chem. Soc. 2022, 144 (1), 18532.

